# Smolt migration in relation to latitude, season and migratory distance

**DOI:** 10.64898/2026.06.20.733518

**Authors:** Madeleine Berry, Benedikte Austad, Kim Aarestrup, Jan G. Davidsen, Marie Nevoux, Carlos M. Alexandre, Sara S. Silva, Jamie R. Stevens, R. Andrew King, Eva B. Thorstad, Johan Höjesjö

**Author notes:** Corresponding author: Madeleine Berry, Department of Biological and Environmental Sciences, University of Gothenburg, 413 90, Gothenburg, Sweden.

## Abstract

Downstream migration and sea entry are periods of high mortality for sea trout smolts and migration timing is a critical aspect of survival. We aimed to investigate migration timing of PIT-tagged sea trout smolts across a latitudinal gradient in five freshwater systems across Europe: Norway, Sweden, Denmark, France and Portugal. In two systems, smolt migration was further examined to assess a) differences in sex, body size and condition between autumn and spring migrants; b) the influence of spatial origin within the stream; and c) relationships between individual body size and migration date. Tagged smolts were not detected migrating in the Portuguese watershed. Spring migration timing differed significantly between watersheds in Norway, Sweden, Denmark and France. Generally, there was a trend of earlier migration at lower latitudes. Autumn vs spring migration was examined in Gudsø-Denmark. Autumn migrants were larger in both length and mass, with no differences were found in sex ratios or body condition. In terms of spatial origin, fish originating from an upstream site were more likely to migrate in the autumn compared to the spring and vice versa. Size dependent migration was found in the Swedish system, Haga å-Sweden, with larger individuals migrating earlier in the spring than smaller individuals. Outward-migrating smolts were also more likely to originate from a downstream site than an upstream site. Overall, these results show both large-scale geographic and fine-scale individual influences on migration timing. Given that climate change may have large impacts on migration patterns in sea trout, understanding variability in migratory patterns across a latitudinal gradient is an important tool for predicting responses to environmental changes.

## Introduction

Migration is an ecological process observed in many species across a wide range of taxa that enables individuals to exploit spatially and temporally variable resources. Salmonids, including the brown trout (*Salmo trutta*), of which the anadromous individuals are commonly known as sea trout, are among the most widely recognized migratory fishes. Anadromous salmonids begin life in freshwater and migrate to sea as juveniles, known as smolts, to take advantage of the richer feeding opportunities in the marine environment, before returning to freshwater to spawn as adults (Nevoux et al., 2019). *Salmo trutta* are an interesting case as they display facultative partial migration; not all individuals undergo anadromous migration, with some remaining in freshwater their entire lives as resident trout (Lobón-Cervía & Sanz, 2018). The decision to migrate is determined at the juvenile stage and reflects a complex interplay of genetic and environmental factors (Ferguson et al., 2019). Aspects such as growth in the preceding autumn (Acolas et al., 2012) and local conditions (Jones et al., 2015) are also important in the decision to migrate.

Timing of migration is a critical determinant of survival (Stich et al., 2015). In sea trout smolts, downstream migration typically occurs in the spring and within a relatively wide temporal migration window of several weeks (Harvey et al., 2020). In sea trout earlier migration is generally associated with higher growth at sea and an increased chance of survival (Jensen et al., 2022). In salmonids, migration is triggered by spring-related changes in photoperiod, temperature and discharge (Aldvén et al., 2015; Zydlewski et al., 2005). Some populations use rising water temperatures during spring as a cue for downstream migration; however, the specific temperature threshold varies among systems. For example, sea trout smolts have been reported as initiating migration at approximately 8°C in a Norwegian system (Haraldstad et al., 2017), and approximately 10°C in a Swedish system (Aldvén et al., 2015). Zydlewski et al., 2005 further proposed that cumulative thermal experience may be more relevant than a single threshold temperature. Although smolt migration is typically associated with spring, there is increasing evidence of an autumn migration period for sea trout and factors such as body size may play a role in determining seasonal migration (Aarestrup et al., 2018; Birnie-Gauvin et al., 2021; Wynne et al., 2023).

Temperature generally increases with decreasing latitude thus southern populations typically experience earlier spring warming and likely an earlier cue in the season to migrate. Consistent with this pattern, migration timing in Atlantic salmon smolts has been linked to latitude (Hvidsten et al., 1998) as well as coho salmon smolts, *Oncorhynchus kisutch*, (Spence & Hall, 2010) with populations at lower latitudes exhibiting earlier migration than those at higher latitudes. With the looming threat of climate change these thermally driven phenological patterns may shift (Schwartz et al., 2006), which could have consequences for survival (Edwards & Richardson, 2004; McCormick et al., 2009). Long-term studies have already documented these shifts in smolt migration (Vehanen et al., 2023), along with effects on survival, particularly in relation to thermal differences between marine and freshwater environments (Kennedy & Crozier, 2010). Because salmonid smolt survival is highest when ocean entry coincides with seasonal peaks in marine productivity driven by plankton blooms, continued climate-driven phenological change may increase the frequency of mismatches between migration timing and optimal marine conditions (Chittenden et al., 2010).

While latitude and temperature shape broad phenological trends, individual characteristics are also of importance. Young fish need to reach a sufficient size as well as accumulate sufficient energy reserves to successfully migrate. Smaller sea trout smolts face higher mortality risk than larger smolts (Dieperink et al., 2001). Recent work also indicates that body condition can differ substantially among migrants from different seasons, with spring migrants being in better condition than autumn migrants suggesting seasonal differences in energetic preparedness in sea trout (Birnie-Gauvin et al., 2021; Winter et al., 2016). Sex-specific differences in migration have also been documented; in many cases the spring smolt run is strongly female-biased with up to 75% of outward-migrating individuals being female (Nevoux et al., 2019) and sex ratios have been found to be similar in autumn and spring migration periods (Birnie-Gauvin et al., 2021).

Migratory distance may likewise influence migration success and factors into smolt mortality, migration timing and subsequent survival (Artero et al., 2023; McCormick et al., 2014; Thorstad et al., 2012). With increased distance comes increased energetic cost as well as increased chance of encountering predators or migration barriers. Therefore, proximity to the marine environment may be an important predictor of successful marine entry, although it is worth noting that there is large variability between sea trout systems, and the trade-offs between long and short migrations may differ substantially between systems. Migratory distance has been linked to seasonal patterns of smolt migration in Atlantic salmon, *Salmo salar*, with autumn migrants most likely to originate from sites closer to the marine environment and spring migrants more likely to originate from further upstream (Ibbotson et al., 2013). Similarly, Jones et al. (2025) showed that distance from the sea influenced the contribution of anadromous mothers to young-of-the-year in a chalk stream system, with fry sampled close to the tidal limit having 100% anadromous mothers, declining to 43% in the headwaters.

Here we investigate timing of migration in sea trout smolts relating to latitude in five freshwater systems across Europe. Additionally, in two of these systems this has been examined further to assess a) relationships between individual body size and date of migration; b) the influence of spatial origin within the stream; and c) differences in sex, body size and condition between autumn and spring migrants. We predict that migration will occur earlier at lower latitudes, that smaller individuals will migrate later within the migration window, and that smolt output will be biased toward individuals originating from downstream sites. Finally, we expect that autumn migrants will be in poorer condition than spring migrants and that sex ratios will be similar between these migration periods with a female bias.

## Methods

### 2.1 Sampling

Five sea trout systems across Europe were selected, covering the Northern and Southern limits of the anadromous populations: Lake Fjærvatnet-Norway, Haga å-Sweden, Gudsø-Denmark, Oir-France and Mouro-Portugal (Fig 1). At each locality an upstream and a downstream site were established separated by < 1 km. In the autumn these sites were electrofished and all juvenile trout caught were anaesthetized, fork length measured, weighed and tagged. The tagging procedure consisted of making a small incision in the abdomen and inserting a 12 mm or 23 mm (depending on fish size) passive integrated transponder (PIT) tag into the body cavity. Each of these tags contains a unique code for individual identification. During sampling at - Haga å-Sweden, Gudsø-Denmark and Fjærvatnet-Norway a small tissue sample was taken, stored in ethanol and sent to a lab for sex determination following the methods of King et al., 2023. After a recovery period, fish were released back into the stream. PIT tag sample sizes can be found in Table 1 and for additional site-specific details see appendix.

**Table 1.** Number of juvenile trout PIT tagged during autumn sampling (Fjærvatnet-Norway 2022, Haga å-Sweden 2021, Gudsø-Denmark 2021, Oir-France 2022, Mouro-Portugal 2023)

| Location | Site | No. Tagged |
| --- | --- | --- |
| Fjærvatnet-Norway | Downstream | 50 |
| Fjærvatnet-Norway | Upstream | 94 |
| Haga å-Sweden | Downstream | 75 |
| Haga å-Sweden | Upstream | 125 |
| Gudsø-Denmark | Downstream | 109 |
| Gudsø-Denmark | Upstream | 92 |
| Oir-France | Downstream | 127 |
| Oir-France | Upstream | 149 |
| Mouro-Portugal | Downstream | 20 |
| Mouro-Portugal | Upstream | 20 |

**Figure 1.**
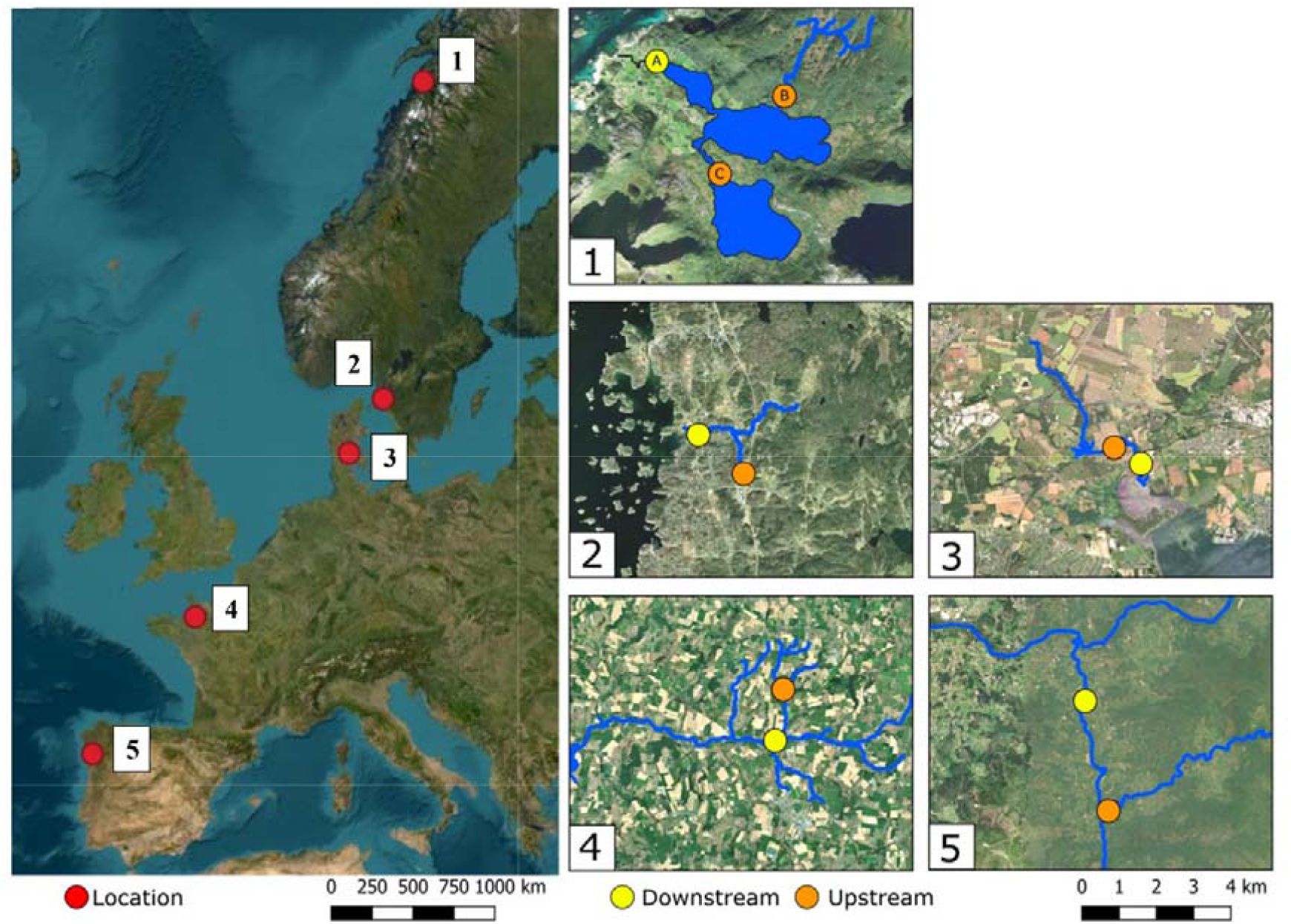
Maps of the 5 study locations (1: Fjærvatnet-Norway, 2: Haga å-Sweden, 3: Gudsø-Denmark, 4: Oir-France and 5: Mouro-Portugal. The left panel shows their positions within Europe. Each stream system is outlined in blue with the various electrofishing sampling sites depicted with yellow for downstream sites and orange for upstream sites (Note: for system 1 site C was not used in this study). Produced using QGIS with ESRI Satellite (ArcGIS/World Imagery) used as a basemap (Scale 1:73296). Adapted from Berry et al., 2025

Individuals were detected during their downstream migration by a combination of smolt traps and stationary PIT tag antennas. At each site, the monitoring period and method differed slightly due to logistical constraints. The monitoring for Fjærvatnet-Norway was conducted using a series of stationary PIT antennas which were operational between 14/04/2023 - 20/06/2023. In Haga å-Sweden a smolt trap was run March - June in 2022, 2023 and 2024. All smolts caught in the smolt trap were scanned for PIT tags and fork length was measured. In Gudsø-Denmark, a stationary PIT tag antenna system runs year-round. In Oir-France, both a smolt trap and stationary PIT antennas system were in place and run year-round. In this study fish detected in both the trap and most downstream antenna were used. In Mouro-Portugal monitoring was conducted using a smolt trap during March – May 2024, no tagged fish were detected migrating during this period; therefore, this system could not be included within our analysis.

### 2.2 Statistical analysis

#### 2.2.1 Autumn vs spring migration

At Gudsø-Denmark, individuals detected passing a stationary PIT antenna at the river mouth were split into autumn migrants, (September – November, n = 29), and spring migrants (February – April, n =27), based on the two assumed peaks in smolt migration in this system (Birnie-Gauvin et al., 2021). Migration season was modelled as a binary response with sex and origin (upstream vs downstream) as predictors using a logistic regression with a binomial error distribution. Differences in size and condition (measured at time of tagging, the autumn prior to migration) were assessed by migration season (autumn vs spring) using separate linear models for condition factor, fork length, and body mass (log-transformed to better fit assumption of normality). Each model included season, sex, and origin as fixed factors and tested with type III sums of squares. Model residuals were examined to verify assumptions of linear regression.

#### 2.2.2 Migration timing across watersheds

In Fjærvatnet-Norway, Haga å-Sweden, Gudsø-Denmark, Oir-France and Mouro-Portugal fish that had been tagged the autumn prior to migration were detected during their outward migration in the spring. Migration timing was analysed using day of the year (DOY). To test whether migration timing differed among the watersheds, a Kruskal–Wallis test was used, as DOY was not normally distributed and sample sizes differed between sites. This was further analysed using pairwise Wilcoxon rank-sum tests with Bonferroni correction to identify which sites differed. To examine whether differences in migration timing were related to latitude, the median DOY for each site was calculated and analysed with a linear regression against the latitude of the sampling location (Fjærvatnet-Norway = 67°, Haga å-Sweden = 57°, Gudsø-Denmark = 55°, Oir-France = 48°).

#### 2.2.3 Size and migration timing

To investigate the relationship between migration timing and body size a linear regression model with fork length against day of the year and year as a categorical factor was made. These data came from all smolts caught in the trap at Haga å-Sweden in the years 2022, 2023 and 2024, regardless of tagging status. Residuals showed heteroskedasticity (Breusch-Pagan test, p < 0.001); therefore, a weighted least squares regression with weights estimated from the relationship between absolute residuals and fitted values of the initial model was used instead. All analyses were conducted in RStudio using the packages: lmtest (Zeileis & Hothorn, 2002), and sandwich (Zeileis et al., 2020). In 2022 and 2023, tagged fish detected from smolt traps (March-June) were analysed to test for differences in recapture probability between fish originating from upstream and downstream. Fisher’s exact two-tailed tests were used, due to small sample sizes, to compare the proportions of recaptured fish between sites for each year and for the combined 2022–2023 dataset. Odds ratios (ORs) and 95% confidence intervals (CIs) were calculated to quantify differences in recapture rate.

## Results

### 3.1 Spring vs. autumn migration (Gudsø-Denmark)

For tagged individuals detected out-migrating at Gudsø-Denmark the migration season (spring/autumn) differed significantly by migratory distance, with individuals originating from the upstream site being more likely to migrate during the autumn than during the spring (Figure 2, Table 2). No difference in sex ratio was detected between autumn and spring migration (Figure 2, Table 2). Both fork length (cm) and body mass (g) differed significantly between migration seasons, with autumn migrants being generally larger at time of tagging in both length and mass than spring migrants (Figure 3, Table 2). However, no difference in condition was detected. Fork length and body mass were also significantly influenced by origin, whereby fish originating from the upstream site were generally larger in both length and mass, whereas sex had no effect on either trait (Table 2).

**Table 2.** Model results testing effects of sex and origin (upstream vs downstream) on migration season (autumn vs spring) and morphological traits of seasonal migrants from Gudsø-Denmark. Migration season was analysed using logistic regression, and condition factor, fork length, and body mass (log-transformed) using linear models with type III sums of squares. Test statistics are z values (logistic model) and F values (linear models). Significant results (p<0,05) are highlighted in bold).

| **Response** | **Predictor** | **Test statistic** | **p-value** |
| --- | --- | --- | --- |
| **Migration season** | Origin | $z = 2.36$ | **0.018** |
| | Sex | $z = 0.69$ | 0.49 |
| **Condition factor (K)** | Season | $F = 0.90$ | 0.35 |
| | Sex | $F = 0.30$ | 0.59 |
| | Origin | $F = 0.63$ | 0.43 |
| **Fork length (cm)** | Season | $F = 8.45$ | **0.0056** |
| | Sex | $F = 1.11$ | 0.30 |
| | Origin | $F = 7.06$ | **0.011** |
| **Mass (g)** | Season | $F = 8.62$ | **0.0051** |
| | Sex | $F = 0.76$ | 0.39 |
| | Origin | $F = 6.70$ | **0.013** |

**Figure 2.**
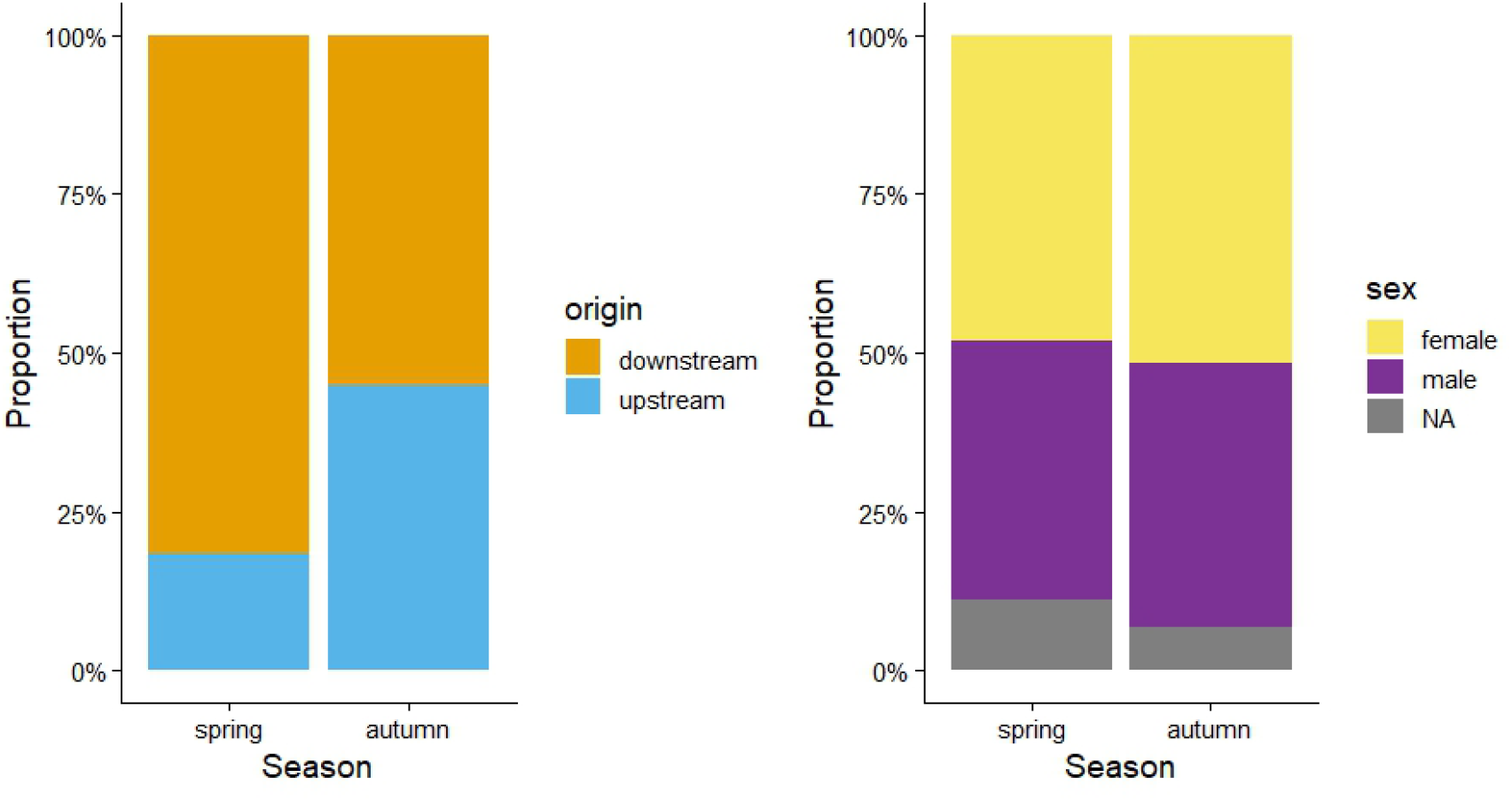
Left: Proportion (%) of autumn and spring migrants that originated from the upstream (blue) and downstream (orange) sites at Gudsø-Denmark. Right: Proportion (%) of autumn and spring migrants that were male (purple), female (yellow), or their sex could not be determined (grey).

**Figure 3.**
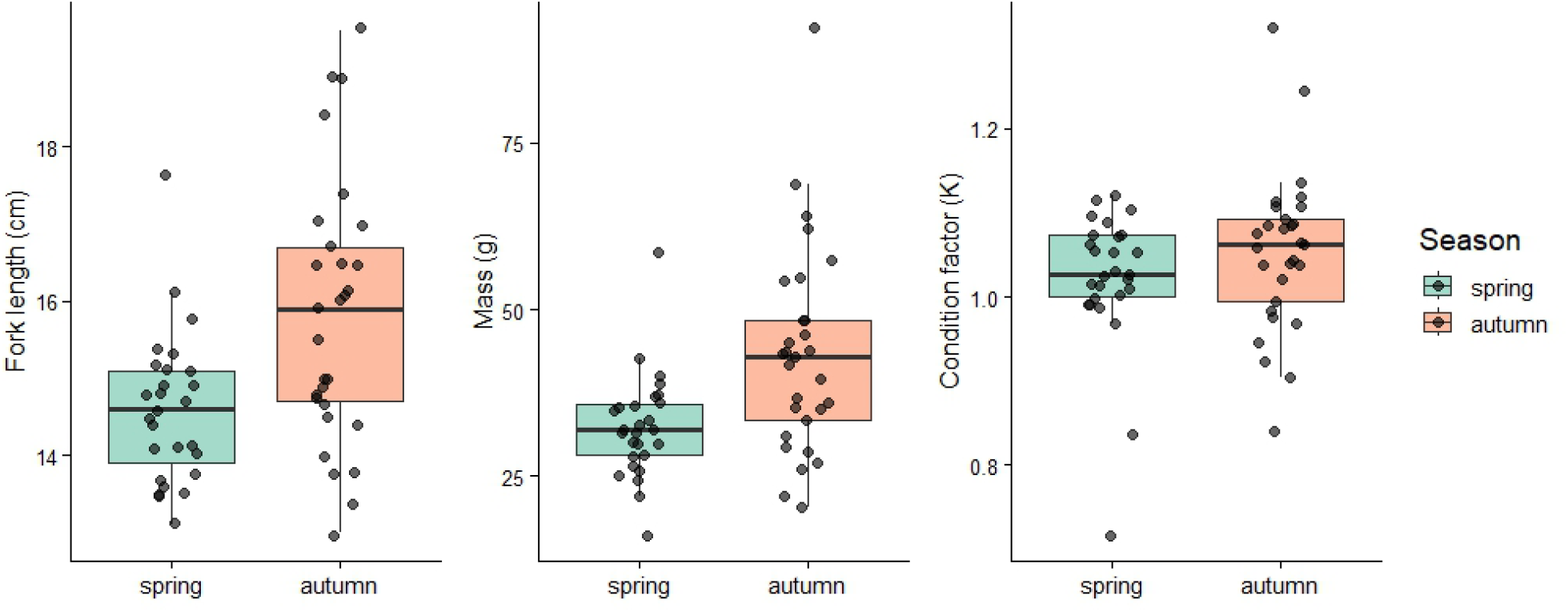
Boxplots showing the differences in fork length (mm) (left), mass (g) (middle) and Fultons condition factor (K) (right) ween autumn (orange) and spring (green) migrants. Boxes include the interquartile range with medians indicated as horizontal lines. Whiskers include values within 1.5 × the interquartile range, and points show individual datapoints.

### 3.2 Migration timing across watersheds

During the spring migration March – July; in Fjærvatnet-Norway, nine individuals were detected migrating, of these three were male and 6 female, 2 originated from upstream and 7 from downstream. In Haga å-Sweden, 13 individuals were detected migrating, 4 originated from upstream and 9 from downstream. In Gudsø-Denmark, 24 individuals were detected migrating, of these 12 were male and 10 female (2 samples could not be determined), 3 originated from upstream and 21 from downstream. In Oir-France, 20 individuals were detected migrating, 11 of which had known origin, 9 originating from upstream and 2 from downstream. When accounting for differences in tagging effort, migration rates differed substantially between upstream and downstream sections. In Fjærvatnet-Norway, Haga å-Sweden, Gudsø-Denmark, downstream fish exhibited higher migration rates (12.0 – 19.3%) than upstream fish (2.1 – 3.3%). In contrast, Oir-France showed the opposite pattern, with a higher migration rate among upstream fish (6.0%) than downstream fish (1.6%). These results suggest watershed specific variation in the relationship between site of origin and migration propensity.

Timing of migration (DOY) differed significantly among watersheds in the different countries during March-July (Kruskal–Wallis test; χ^2^ = 35.71, df = 3, p < 0.001) (Figure 4). Post-hoc pairwise Wilcoxon tests with Bonferroni correction showed that all country pairs differed significantly (p < 0.05) except for Gudsø-Denmark and Oir-France, which did not differ (p = 1.0). Median migration dates occurred earliest in Gudsø-Denmark (4th April) and Oir-France (8th April), followed by Haga å-Sweden (24th April), and Fjærvatnet-Norway (11th June). The central 50% of migration occurred between 3rd March – 18th April in Gudsø-Denmark, 24th March – 13th April in Oir-France, 23rd April – 15th May in Haga å-Sweden, and 21st May – 18th June in Fjærvatnet-Norway. The relationship between latitude and median DOY was assessed with a linear regression and showed a marginal effect of latitude (β = 3.59, SE = 1.20, R^2^ = 0.82, p = 0.096), indicating migration occurred approximately 3.6 days later per degree of latitude. These results indicate that smolt migration timing varies significantly among the studied watersheds in the different countries and this was associated with latitude.

**Figure 4.**
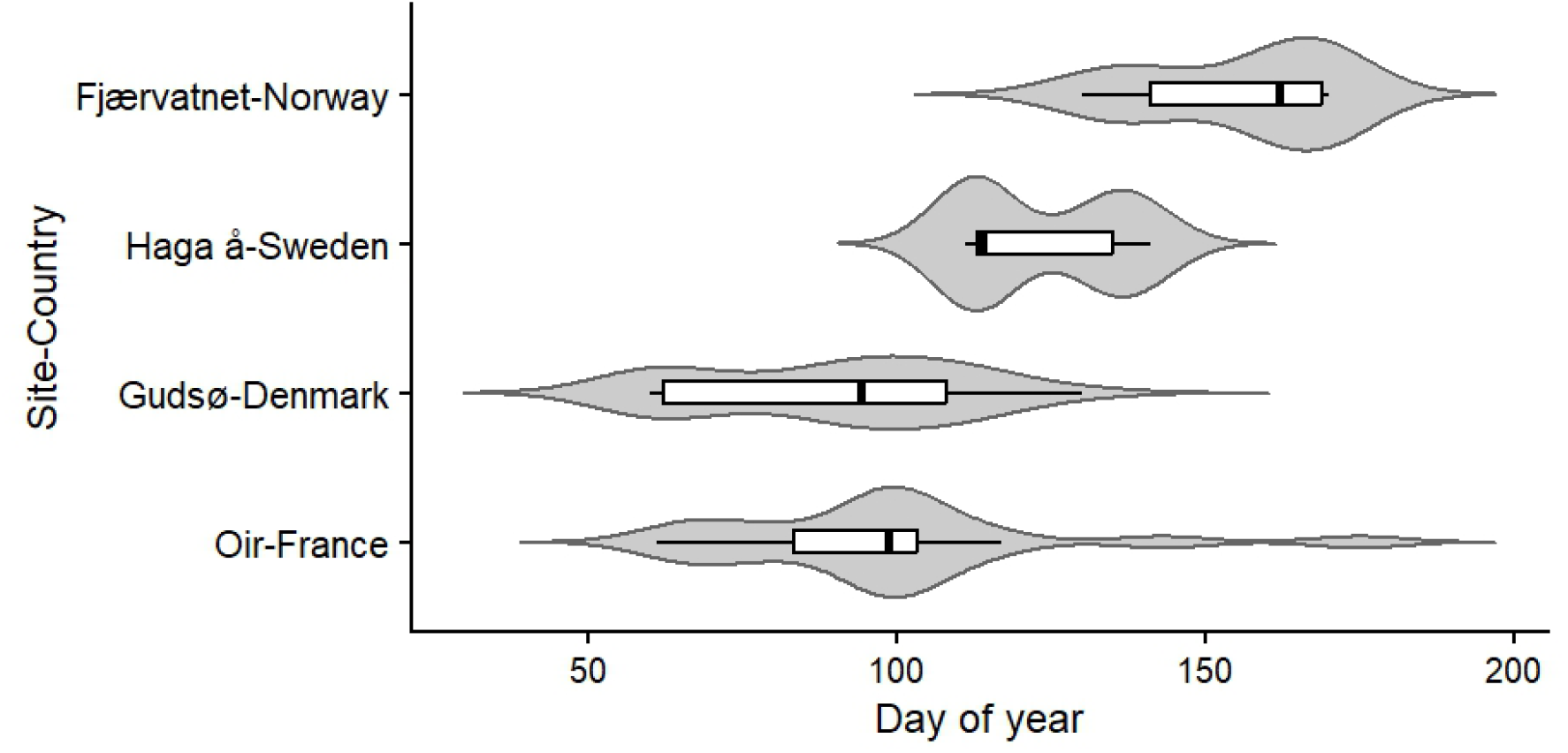
Distribution of migration timing (day of year) for watersheds in each country between March and July, Fjærvatnet-Norway = 2023, Haga å-Sweden = 2022, Gudsø-Denmark = 2022, Oir-France = 2023. Violin plots show the distribution of observations, and boxplots show medians and interquartile ranges with whiskers extending to the most extreme values within 1.5 x the interquartile range.

### 3.3 Size and migration timing (Haga å-Sweden)

Using trap data from 2022 – 2024, a weighted least squares regression of fish length at time of migration against day of year and year as a factor (using 2022 as a baseline) showed a negative relationship between day-of-year and length (t = −15.3, SE = 0.03, p < 0.001), indicating larger smolts migrated earlier in the season than smaller smolts (Figure 5). Additionally, fish were on average larger in 2023 than in 2022 (t = 2.3, SE = 21.1, p = 0.02) and the slope was marginally steeper (t = −1.9, SE = 0.2, p = 0.06), whereas 2024 did not significantly differ from 2022 in average length (t = 0.2, SE = 56.4, p = 0.81) or slope (t = −0.2, SE = 0.5, p = 0.87).

**Figure 5.**
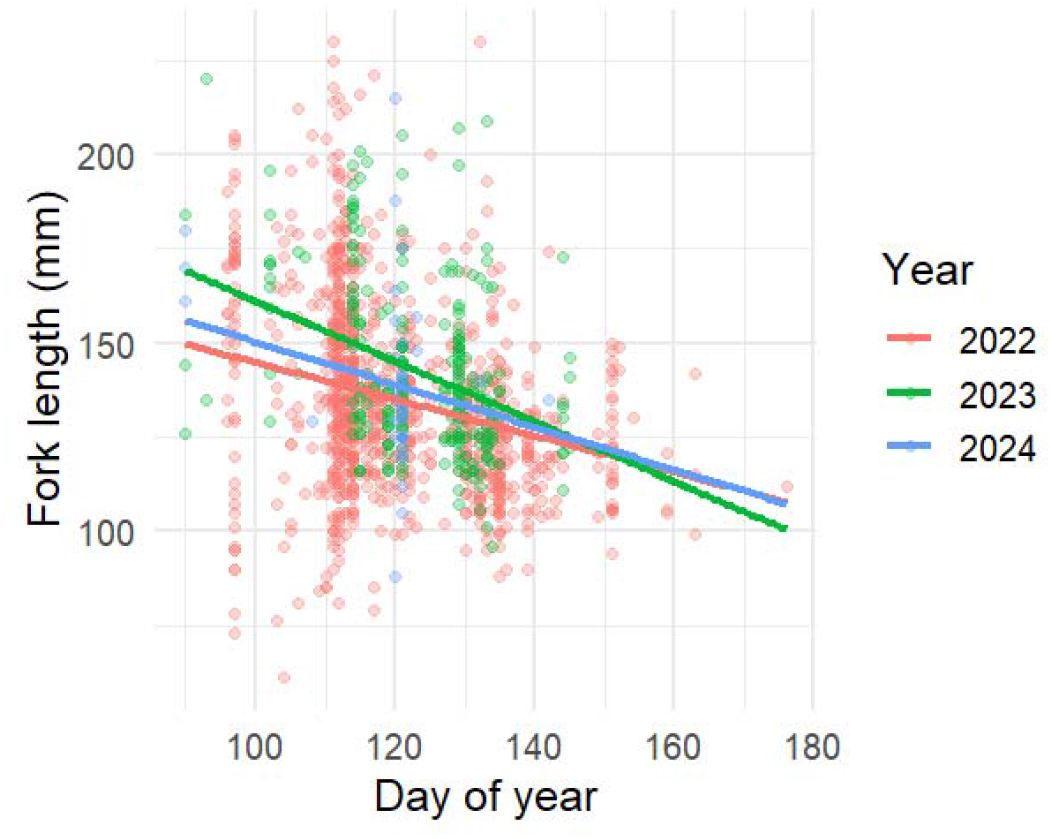
Raw data of fork length and migration day with the fitted WLS regression lines for 2022 (red), 2023 (green) and 2024 (blue) for smolts caught in a fish trap migrating during March – June at Haga å-Sweden.

When data from 2022 and 2023 were pooled, recapture rate was significantly higher (Fisher’s exact test; odds ratio = 0.28, 95 % CI = 0.11-0.71, p = 0.004) for fish originating from downstream (22 %) than those originating from upstream (7 %). In 2022, recapture rate did not differ significantly between fish originating from upstream (7 %) and downstream (14 %) sites (Fisher’s exact test; odds ratio = 0.44, 95 % CI = 0.09-2.84, p = 0.37). In 2023, fish originating from downstream had a significantly higher recapture rate (27 %) than those originating from upstream (7 %) (Fisher’s exact test; odds ratio = 0.23, 95 % CI = 0.06-0.86, p = 0.015). In total,124 individuals were sampled for sex determination when they were caught in the smolt trap, of which 52 were male (42%) and 72 female (58%).

## Discussion

Smolt migration represents a key transitional phase in the sea trout lifecycle linking freshwater and marine environments. Understanding variability in this phase is critical as it is a period of high mortality and is sensitive to environmental changes. The results of this study demonstrate the variability in downstream migratory patterns; specifically in migration timing and individual characteristics of migrants. Our results suggested a trend that more northern populations migrate later than more southern populations, although this inference is based on a limited number of sites (n = 4) and small sample sizes, the relationship was strong (R^2^ = 0.82), indicating that latitude may be an important driver of smolt migration timing. Large-scale geographic variation between watersheds in the onset of the migration window suggests possible links with latitude-related climatic variables which is also reflected in the wider literature (Figure 6). Variation was also captured in migration propensity and site of origin, with individuals originating from downstream sections more likely to be detected migrating than those originating from upstream sites in the Scandinavian watersheds and the reverse in the French watershed. It would be valuable to continue these types of timing studies with more sites across a broader range of latitudes. Additionally, detection was determined here using both fish traps and stationary PIT antennas due to logistical constraints, but it is worth bearing in mind there is likely variation in detectability between these methods. The fate of tagged individuals not detected is unknown and some may have migrated and simply not been detected either due to passing a trap without being caught or inefficient stationary PIT antennas.

**Figure 6.**
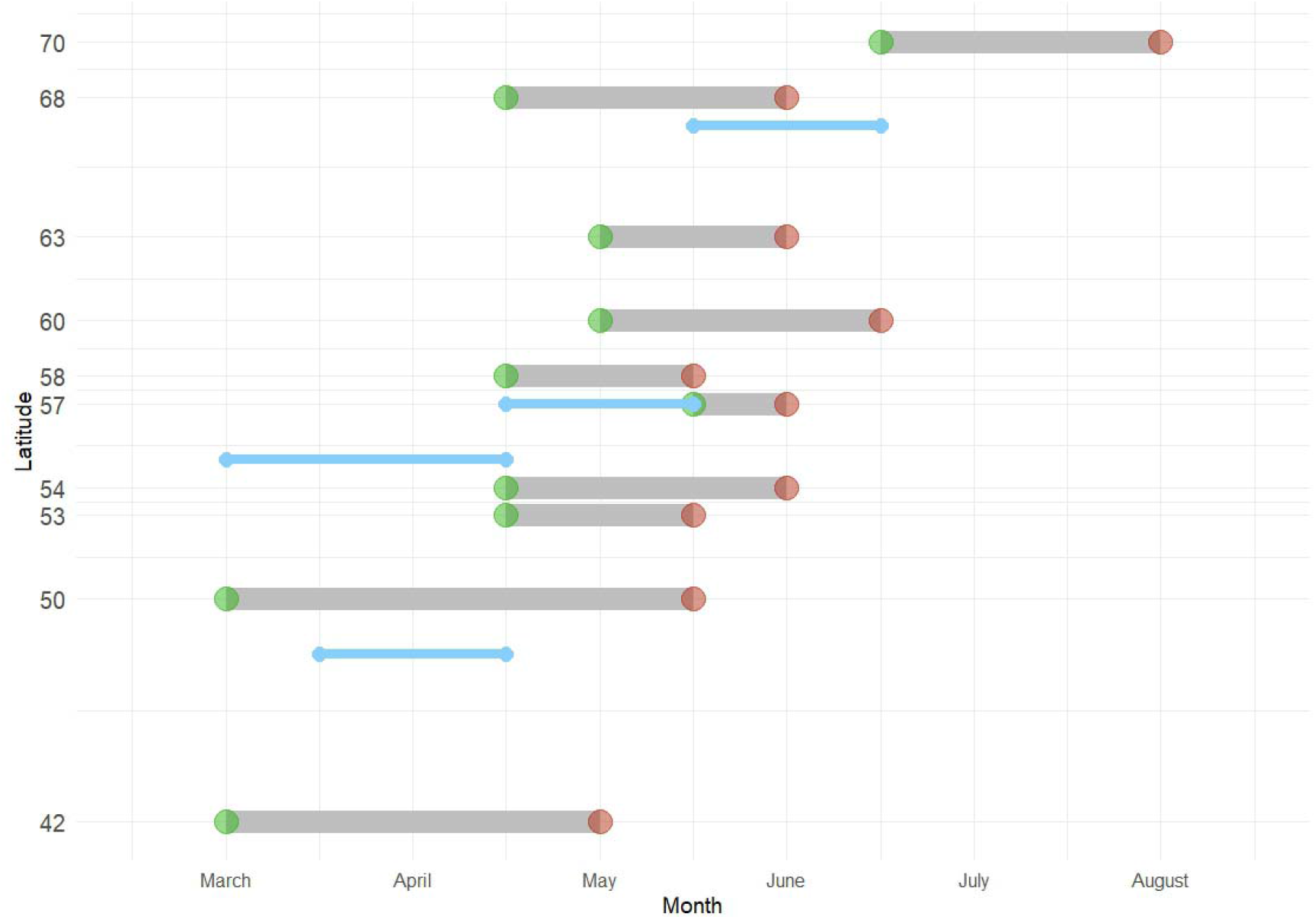
Lollipop chart showing spring migration periods reported in literature across different latitudes approximated to the nearest two weeks and generally representing the central portion of the smolt run although level of detail differed between sources. Green dots represent start times and red dots end times. Sources from top down: Jensen et al. 2012, Sjursen et al. 2021, Hembrel et al. 2001, Harvey et al. 2020, Bohlin et al. 1993, Pratten & Shearer 1983, de Eyto et al. 2002, Gargan et al. 2016, Euzenat et al. 2007, Cabellero et al. 2007. Migration periods from the present study are overlayed in blue, central migration windows were defined as the 25–75% quantile range and rounded to the nearest half-month for visualization.

No tagged individuals were detected migrating at Mouro-Portugal, preventing its inclusion in the statistical analyses. However, this result remains noteworthy given that the site was selected to represent the southern extent of the study’s latitudinal gradient as this is the southern limit of the sea trout native range. The Minho basin where this site is located is the area of Portugal where sea trout is the most abundant. Several explanations may account for the absence of detections aside from the small sample size of tagged fish. First, the proportion of anadromous individuals in this population may be low, with most fish adopting a resident freshwater life-history strategy. Second, migratory individuals may use alternative migration strategies by migrating outside of the spring migration window. The smolt trap operated primarily during the expected spring migration window (March–May), meaning individuals migrating outside of this window are not detected. Finally, mortality during the time between tagging and downstream passage cannot be excluded. Further monitoring across a broader temporal window, particularly using permanent PIT antenna arrays, would help distinguish among these hypotheses. The absence of detected migrants may also be relevant in the context of environmental constraints operating near the southern edge of the species’ distribution. Southern populations of sea trout are expected to experience warmer temperatures, reduced summer flows, and increased hydrological variability compared to more northern populations, all of which may alter the costs and benefits of anadromy. With environmental conditions likely to shift under climate change in the future we may see shifts in migration timings as well as the proportion of anadromy in a population. The persistence of anadromous sea trout at the southern margin of the species’ range may depend on behavioural flexibility and the availability of suitable migration opportunities.

In Gudsø-Denmark, where autumn migration is common there were differences in body size between autumn and spring migrants. Body size and condition were measured at the time of tagging in September, with autumn migration occurring shortly thereafter and spring migration the following year. Autumn migrants were larger in both length and mass than spring migrants. This could be related to age-structuring in migration however age has not been determined for these individuals. No difference in body condition was found between autumn and spring migrants, which contrasts with previous findings from the same system, where autumn migrants were reported to be in poorer condition than spring migrants (Birnie-Gauvin et al., 2021). The size difference demonstrated here could suggest that individuals that have already reached sufficient size for migration may opt for migrating during the autumn rather than waiting and overwintering in the stream to migrate in the spring. In both autumn and spring migrants the sex ratio was approximately 50%, which differs from the assumption that typically there is a bias towards females (Nevoux et al., 2019). Additionally, individuals originating from the upstream site were more likely to migrate in the autumn than the spring and vice versa. Investigating whether origin along the river continuum predicts migration season in other systems would be important for further understanding migration timing variation.

In Haga å-Sweden larger individuals tended to migrate earlier than smaller individuals across three years. This pattern suggests migration timing has an element of size-dependence in this system. Smaller individuals may not have reached an optimal size or accumulated sufficient energy reserves early in the migration season and may incur higher mortality if they migrate prematurely and therefore delay migration. Consequently, later migration may represent a compensatory strategy for smaller or slower-growing individuals. These patterns could also be an indication of migration timing separated by age class with older fish migrating before younger fish, however age was not measured here. In this population, it has previously been shown that individuals from the downstream site are larger than those from the upstream site (Berry et al., 2024), and perhaps this spatial sorting of size could play a role in variation in migration timing. Gillson et al., 2025 found the opposite pattern in sea trout smolts with larger individuals migrating later in the season, suggesting the relationship between size and migration timing varies among populations. Differences in local environmental conditions and growth opportunities may influence the relative advantages of early versus late migration for individuals of different sizes.

The distribution of juveniles within the freshwater system largely reflects the location of parental spawning and differences between upstream and downstream populations can be due to a combination of genetics, maternal effects and environmental effects. Such differences have been observed in traits such as behaviour (Berry et al., 2025; Kristensen & Closs, 2008), stress-related responses (Berry et al., 2024) and migratory strategy (Olsson & Greenberg, 2004). Here, it has been shown that migratory distance may have consequences for migration behaviours and potentially life history strategy. In three systems, individuals confirmed as successful spring outmigrants were more likely to be from the downstream site than the upstream site. This could be the result of a multitude of factors: downstream individuals may be more likely to adopt anadromy compared to upstream individuals who may favour residency, individuals from upstream may experience greater mortality during their downstream migration and thus fewer survive to migrate out, or individuals originating from upstream could migrate at other times outside of the typical migration window. In Gudsø-Denmark, individuals from the upstream site were more likely to migrate during the autumn migration period than the spring migration period, and perhaps a similar mechanism may be functioning in other systems causing the lower number of upstream migrants compared to downstream found during the spring. These findings are consistent with the idea that spatial structuring within freshwater systems can strongly influence migratory patterns, with implications for how different parts of a catchment contribute to smolt production and temporal variation in migration.

Given the potential for climate change to alter temperature and precipitation conditions there is the potential for environmental triggers for migration to shift. Increases in temperatures may shift the migration peak earlier, while changes in the timing of heavy rainfall could further alter migration timing. While climate change is inevitable, the degree to which brown trout and sea trout, in particular, will be affected remains unknown. Here we have shown some of the scope of migratory patterns and timing, providing important insights into the adaptive capacity of sea trout populations. Climate change is not the sole pressure sea trout face and mitigation measures for other risks such as habitat fragmentation need to be addressed to maintain healthy populations with access to the sea. Restoration of degraded systems and opening of migration routes offers a chance to boost numbers of out-migrating sea trout and buffer populations against losses.

## Acknowledgements

This work was funded by a grant from FORMAS (grant no: 2020-01349). The data collection program on salmonids on the Oir River was carried out by Ecological Research Observatory on Diadromous Fish in coastal streams by INRAE 1036 U3E (ORE DiaPFC 2024).

## Data availability

Data can be found at https://doi.org/10.5281/zenodo.20843935

## Appendix

Additional sampling information (adapted from Berry et al., 2025).

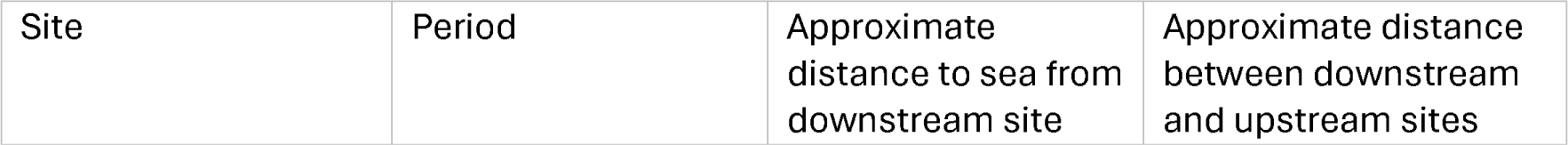
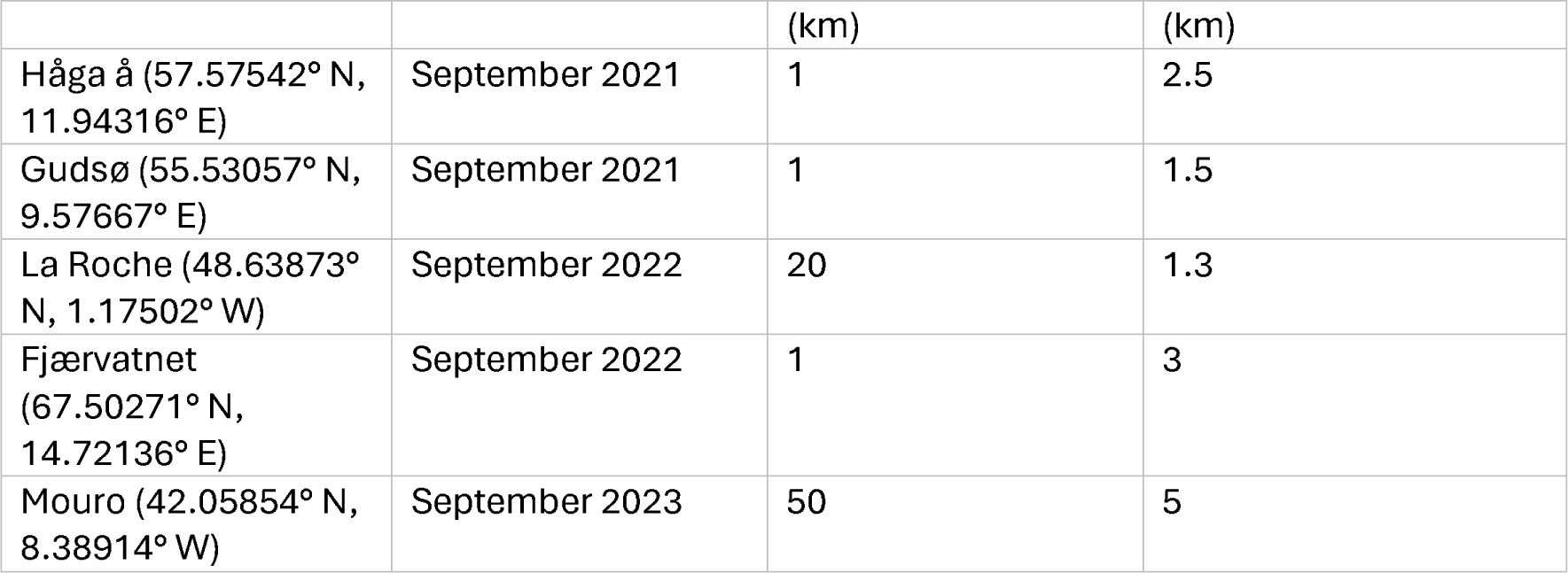

| Site | Period | Approximate distance to sea from downstream site | Approximate distance between downstream and upstream sites |
| --- | --- | --- | --- |

|  |  | (km) | (km) |
| --- | --- | --- | --- |
| Håga å (57.57542° N, 11.94316° E) | September 2021 | 1 | 2.5 |
| Gudsø (55.53057° N, 9.57667° E) | September 2021 | 1 | 1.5 |
| La Roche (48.63873° N, 1.17502° W) | September 2022 | 20 | 1.3 |
| Fjærvatnet (67.50271° N, 14.72136° E) | September 2022 | 1 | 3 |
| Mouro (42.05854° N, 8.38914° W) | September 2023 | 50 | 5 |

## References

Aarestrup, K., Birnie-Gauvin, K., & Larsen, M. H. (2018). Another paradigm lost? Autumn downstream migration of juvenile brown trout: Evidence for a presmolt migration. Ecology of Freshwater Fish, 27(1), 513–516. 10.1111/eff.12335

Acolas, M. L., Labonne, J., Baglinière, J. L., & Roussel, J. M. (2012). The role of body size versus growth on the decision to migrate: A case study with Salmo trutta. Naturwissenschaften, 99(1), 11–21. 10.1007/s00114-011-0861-5

Aldvén, D., Degerman, E., & Höjesjö, J. (2015). Environmental cues and downstream migration of anadromous brown trout (Salmo trutta) and Atlantic salmon (Salmo salar) smolts. 20, 10.

Artero, C., Gregory, S., Beaumont, W., Josset, Q., Jeannot, N., Cole, A., Lamireau, L., Réveillac, E., & Lauridsen, R. (2023). Survival of Atlantic salmon and sea trout smolts in transitional waters. Marine Ecology Progress Series, 709, 91–108. 10.3354/meps14278

Berry, M., Davidsen, J. G., Nevoux, M., Aarestrup, K., Alexandre, C. M., Silva, S. S., Thorén, A., Engstöm, A., Ahvenainen, M., & Höjesjö, J. (2025). Do offspring characteristics reflect parental migration variation? Journal of Fish Biology, jfb.70247.10.1111/jfb.70247

Berry, M., Zena, L. A., Roques, J. A., Sandblom, E., Thorstad, E. B., & Höjesjö, J. (2024). Local variation in stress response of juvenile anadromous brown trout, Salmo trutta. Ecology and Evolution, 14(6), e11526.

Birnie-Gauvin, K., Larsen, M. H., & Aarestrup, K. (2021). Energetic state and the continuum of migratory tactics in brown trout (Salmo trutta). Canadian Journal of Fisheries and Aquatic Sciences, 78(10), 1435–1443. 10.1139/cjfas-2021-0025

Bohlin, T., Dellefors, C., & Faremo, U. (1993). Timing of Sea-Run Brown Trout (Salmo trutta) Smolt Migration: Effects of Climatic Variation. Canadian Journal of Fisheries and Aquatic Sciences, 50(6), 1132–1136. 10.1139/f93-128

Caballero, P., Cobo, F., & Gonzlez, M. A. (2007). Life History of a Sea Trout (Salmo trutta L.) Population from the North-West Iberian Peninsula (River Ulla, Galicia, Spain). In G. Harris & N. Milner (Eds), Sea Trout: Biology, Conservation and Management (pp. 234–247). Blackwell Publishing Ltd. 10.1002/9780470996027.ch16

Chittenden, C. M., Jensen, J. L. A., Ewart, D., Anderson, S., Balfry, S., Downey, E., Eaves, A., Saksida, S., Smith, B., Vincent, S., Welch, D., & McKinley, R. S. (2010). Recent Salmon Declines: A Result of Lost Feeding Opportunities Due to Bad Timing? PLoS ONE, 5(8), e12423. 10.1371/journal.pone.0012423

De Eyto, E., Kelly, S., Rogan, G., French, A., Cooney, J., Murphy, M., Nixon, P., Hughes, P., Sweeney, D., McGinnity, P., Dillane, M., & Poole, R. (2022). Decadal Trends in the Migration Phenology of Diadromous Fishes Native to the Burrishoole Catchment, Ireland. Frontiers in Ecology and Evolution, 10, 915854. 10.3389/fevo.2022.915854

Dieperink, C., Pedersen, S., & Pedersen, M. I. (2001). Estuarine predation on radiotagged wild and domesticated sea trout (Salmo trutta L.) smolts: Estuarine predation on smolts. Ecology of Freshwater Fish, 10(3), 177–183. 10.1034/j.1600-0633.2001.100307.x

Edwards, M., & Richardson, A. J. (2004). Impact of climate change on marine pelagic phenology and trophic mismatch. Nature, 430(7002), 881–884. 10.1038/nature02808

Euzenat, G., Fournel, F., & Richard, A. (1999). Sea Trout (Salmo trutta L.) in Normandy and Picardy. In Biology and ecology of the brown and sea trout (pp. 175–203).

Ferguson, A., Reed, T. E., Cross, T. F., McGinnity, P., & Prodöhl, P. A. (2019). Anadromy, potamodromy and residency in brown trout Salmo trutta: The role of genes and the environment. Journal of Fish Biology, 95(3), 692–718. 10.1111/jfb.14005

Gargan, P., Kelly, F., Shepard, S., & Whelan, K. (2016). Temporal variation in sea trout Salmo trutta life history traits in the Erriff River, western Ireland. Aquaculture Environment Interactions, 8, 675–689. 10.3354/aei00211

Gillson, J. P., Blackwell, R. E., Gregory, S. D., Marsh, J. E., Bašić, T., Elliott, S. A., … & Lauridsen, R. B. (2025). Do the biological characteristics of trout (Salmo trutta) smolts influence their spring migration timing and maiden marine sojourn duration?. Journal of Fish Biology.

Haraldstad, T., Kroglund, F., Kristensen, T., Jonsson, B., & Haugen, T. O. (2017). Diel migration pattern of Atlantic salmon (Salmo salar) and sea trout (Salmo trutta) smolts: An assessment of environmental cues. Ecology of Freshwater Fish, 26(4), 541–551. 10.1111/eff.12298

Harvey, A. C., Glover, K. A., Wennevik, V., & Skaala, Ø. (2020). Atlantic salmon and sea trout display synchronised smolt migration relative to linked environmental cues. Scientific Reports, 10(1), 3529. 10.1038/s41598-020-60588-0

Hembrel, B., Arnekleiv, J. V., & L’Abee-Lund, J. H. (2001). Effects of water discharge and temperature on the seaward migration of anadromous brown trout, Salmo trutta, smolts. Ecology of Freshwater Fish, 10(1), 61–64.

Hvidsten, N. A., Heggberget, T. G., & Jensen, A. J. (1998). Sea water temperatures at Atlantic salmon smolt enterance. Nordic Journal of Freshwater Research, 79–86.

Ibbotson, A. T., Riley, W. D., Beaumont, W. R. C., Cook, A. C., Ives, M. J., Pinder, A. C., & Scott, L. J. (2013). The source of autumn and spring downstream migrating juvenile A tlantic salmon in a small lowland river. Ecology of Freshwater Fish, 22(1), 73–81. 10.1111/eff.12003

Jensen, A. J., Diserud, O. H., Finstad, B., Fiske, P., & Thorstad, E. B. (2022). Early□season brown trout (Salmo trutta) migrants grow and survive better at sea. Journal of Fish Biology, 100(6), 1419–1431. 10.1111/jfb.15052

Jensen, A. J., Finstad, B., Fiske, P., Hvidsten, N. A., Rikardsen, A. H., & Saksgård, L. (2012). Timing of smolt migration in sympatric populations of Atlantic salmon (Salmo salar), brown trout (Salmo trutta), and Arctic char (Salvelinus alpinus). Canadian Journal of Fisheries and Aquatic Sciences, 69(4), 711–723. 10.1139/f2012-005

Jones, D. A., Bergman, E., & Greenberg, L. (2015). Food availability in spring affects smolting in brown trout (Salmo trutta). Canadian Journal of Fisheries and Aquatic Sciences, 72(11), 1694–1699. 10.1139/cjfas-2015-0106

Jones, J. I., R. A. King, A. M. Sturrock, H. Wei, J. R. Stevens, & R. B. Lauridsen. 2025. Anadromous brown trout (Salmo trutta) contribute disproportionally to recruitment: Insights from genomics and multi-tissue stable isotope chemistry. Freshwater Biology 70, no. 8: e70078. 10.1111/fwb.70078.

King, R. A., Toms, S., & Stevens, J. R. (2023). Evaluating the importance of accurate sex ratios on egg deposition targets and conservation limit compliance for Atlantic salmon (Salmo salar L.) in the River Tamar, south-west England. Fisheries Management and Ecology, 30, 161–170. 10.1111/fme.12609

Kennedy, R. J., & Crozier, W. W. (2010). Evidence of changing migratory patterns of wild Atlantic salmon Salmo salar smolts in the River Bush, Northern Ireland, and possible associations with climate change. Journal of Fish Biology, 76(7), 1786–1805. 10.1111/j.1095-8649.2010.02617.x

Kristensen, E. A., & Closs, G. P. (2008). Variation in growth and aggression of juvenile brown trout (Salmo trutta) from upstream and downstream reaches of the same river. Ecology of Freshwater Fish, 17(1), 130–135. 10.1111/j.1600-0633.2007.00266.x

Lobón-Cervía, J., & Sanz, N. (2018). Brown trout: Biology, ecology and management. Wiley.

McCormick, S. D., Haro, A., Lerner, D. T., O’Dea, M. F., & Regish, A. M. (2014). Migratory patterns of hatchery and stream□reared Atlantic salmon Salmo salar smolts in the Connecticut River,

U.S.A. Journal of Fish Biology, 85(4), 1005–1022. 10.1111/jfb.12532

McCormick, S. D., Lerner, D. T., Monette, M. Y., Nieves-Puigdoller, K., Kelly, J. T., & Björnsson, B. T. (2009). Taking It with You When You Go: How Perturbations to the Freshwater Environment, Including Temperature, Dams, and Contaminants, Affect Marine Survival of Salmon. American Fisheries Society Symposium, 69, 195–214.

Nevoux, M., Finstad, B., Davidsen, J. G., Finlay, R., Josset, Q., Poole, R., Höjesjö, J., Aarestrup, K., Persson, L., Tolvanen, O., & Jonsson, B. (2019). Environmental influences on life history strategies in partially anadromous brown trout (Salmo trutta, Salmonidae). Fish and Fisheries, 20(6), 1051–1082. 10.1111/faf.12396

Olsson, I. C., & Greenberg, L. A. (2004). Partial migration in a landlocked brown trout population. Journal of Fish Biology, 65(1), 106–121. 10.1111/j.0022-1112.2004.00430.x

Pratten, D. J., & Shearer, W. M. (1983). The migrations of North Esk sea trout. Aquaculture Research, 14(3), 99–113.

Schwartz, M. D., Ahas, R., & Aasa, A. (2006). Onset of spring starting earlier across the Northern Hemisphere. Global Change Biology, 12(2), 343–351. 10.1111/j.1365-2486.2005.01097.x

Sjursen, A. D., Friis, M. E. L., Rønning, L., & Davidsen, J. G. (2021). Overvåkning av anadrome laksefisk i Fjærevassdraget, Nordland. Resultater fra videoovervåkningen i 2020. NTNU Vitenskapsmuseet naturhistorisk rapport.

Spence, B. C., & Hall, J. D. (2010). Spatiotemporal patterns in migration timing of coho salmon (Oncorhynchus kisutch) smolts in North America. Canadian Journal of Fisheries and Aquatic Sciences, 67(8), 1316–1334. 10.1139/F10-060

Stich, D. S., Zydlewski, G. B., Kocik, J. F., & Zydlewski, J. D. (2015). Linking Behavior, Physiology, and Survival of Atlantic Salmon Smolts During Estuary Migration. Marine and Coastal Fisheries, 7(1), 68–86. 10.1080/19425120.2015.1007185

Thorstad, E. B., Whoriskey, F., Uglem, I., Moore, A., Rikardsen, A. H., & Finstad, B. (2012). A critical life stage of the Atlantic salmon Salmo salar: Behaviour and survival during the smolt and initial post□smolt migration. Journal of Fish Biology, 81(2), 500–542. 10.1111/j.1095-8649.2012.03370.x

Vehanen, T., Sutela, T., & Huusko, A. (2023). Potential Impact of Climate Change on Salmonid Smolt Ecology. Fishes, 8(7), 382. 10.3390/fishes8070382

Winter, E. R., Tummers, J. S., Aarestrup, K., Baktoft, H., & Lucas, M. C. (2016). Investigating the phenology of seaward migration of juvenile brown trout (Salmo trutta) in two European populations. Hydrobiologia, 775(1), 139–151. 10.1007/s10750-016-2720-z

Wynne, R., Kaufmann, J., Coughlan, J., Phillips, Karl. P., Waters, C., Finlay, R. W., Rogan, G., Poole, R., McGinnity, P., & Reed, T. E. (2023). Autumn outmigrants in brown trout (Salmo trutta) are not a demographic dead□end. Journal of Fish Biology, 102(6), 1327–1339. 10.1111/jfb.15377

Zeileis A, Köll S, Graham N (2020). “Various Versatile Variances: An Object-Oriented Implementation of Clustered Covariances in R.” Journal of Statistical Software, 95(1), 1–36. doi:10.18637/jss.v095.i01.

Zeileis A, Hothorn T (2002). “Diagnostic Checking in Regression Relationships.” R News, 2(3), 7–10. https://CRAN.R-project.org/doc/Rnews/.

Zydlewski, G. B., Haro, A., & McCormick, S. D. (2005). Evidence for cumulative temperature as an initiating and terminating factor in downstream migratory behavior of Atlantic salmon (Salmo salar) smolts. Canadian Journal of Fisheries and Aquatic Sciences, 62(1), 68–78. 10.1139/f04-179

